# TET-catalyzed 5-carboxylcytosine promotes CTCF binding to suboptimal sequences genome-wide

**DOI:** 10.1101/480525

**Authors:** Kyster K. Nanan, David M. Sturgill, Maria F. Prigge, Morgan Thenoz, Allissa A. Dillman, Mariana D. Mandler, Shalini Oberdoerffer

## Abstract

The mechanisms supporting dynamic regulation of CTCF binding sites remain poorly understood. Here we describe the TET-catalyzed 5-methylcytosine derivative, 5-carboxylcytosine (5caC) as a factor driving new CTCF binding within genomic DNA. Through a combination of *in vivo* and *in vitro* approaches, we reveal that 5caC generally strengthens CTCF association with DNA and facilitates binding to suboptimal sequences. Dramatically, profiling of CTCF binding in a cellular model that accumulates genomic 5caC identified ∼13,000 new CTCF sites. The new sites were enriched for overlapping 5caC and were marked by an overall reduction in CTCF motif strength. As CTCF has multiple roles in gene expression, these findings have wide-reaching implications and point to induced 5caC as a potential mechanism to achieve differential CTCF binding in cells.

## INTRODUCTION

CTCF is an 11 zinc-finger DNA-binding protein that regulates multiple critical genomic functions, including promoting long-range interactions between distal regions of the genome and insulating areas of active transcription from inactive regions (Phillips and Corces, 2009). Profiling of CTCF binding in human cells suggests tens of thousands of binding sites, more than half of which display tissue specificity (Chen et al., 2012; The ENCODE Project Consortium, 2012; Wang et al., 2012). Regulation of CTCF binding at variable locations is largely achieved through dynamic DNA methylation, wherein overlapping 5-methylcytosine (5mC) inhibits CTCF association with DNA (Bell and Felsenfeld, 2000; Hark et al., 2000; Maurano et al., 2015; Shukla et al., 2011; The ENCODE Project Consortium, 2012; Wang et al., 2012). Amongst the described functions for variable CTCF sites, we identified a role in alternative pre-mRNA splicing (Shukla et al., 2011). CTCF binding within actively transcribed genes transiently obstructs RNA polymerase II (pol II) elongation, thereby kinetically favoring spliceosome assembly at weak upstream splice sites (Shukla et al., 2011). In contrast, inhibition of CTCF binding through overlapping 5mC shifts splicing to competing downstream sites through loss of pol II pausing (Shukla et al., 2011). However, the mechanisms that dynamically regulate CTCF exchange in alternative splicing and other tissue-specific activities remained unknown.

We recently determined that the alpha-ketoglutarate dependent dioxygenases, TET1 and TET2 support CTCF function in splicing regulation through antagonizing overlapping 5-methylcytosine (5mC) at CTCF binding sites (Marina et al., 2016). The TET proteins catalyze active DNA demethylation through successive oxidation of 5mC to 5-hydroxymethycytosine (5hmC), 5-formylcytosine (5fC) and 5-carboxycytosine (5caC) (Ito et al., 2011; Tahiliani et al., 2009). In the final step in the demethylation pathway, 5caC is converted to cytosine through the base excision repair factor thymine DNA glycosylase (TDG) (He et al., 2011; Maiti and Drohat, 2011). In contrast, reduced TET activity results in increased 5mC at CTCF binding sites and associated exclusion of dependent upstream exons from spliced mRNA due to CTCF eviction (Marina et al., 2016). Curiously, the underlying DNA at these splicing-associated CTCF sites was not fully unmethylated but was rather marked by a steady level of 5caC (Marina et al., 2016). While 5caC levels in genomic DNA are generally low, we readily detected the oxidized derivative within CTCF sites in actively dividing primary peripheral lymphocytes (Marina et al., 2016). Biochemical characterization confirmed CTCF interaction with 5caC-containing DNA *in vitro* that was, unexpectedly, enhanced as compared to unmethylated DNA (Marina et al., 2016). However, the significance to CTCF association with 5caC *in vivo* remained unclear.

Here we directly examine whether and how 5caC influences CTCF binding in cells. Through utilizing a cellular system that accumulates 5caC within genomic DNA (Cortazar et al., 2011), we observe a dramatic expansion in locations of CTCF binding, genome-wide. Characterization of CTCF binding at these *de novo* sites revealed unique features, including an enrichment for overlapping 5caC and loosening of the consensus CTCF binding motif. Likewise, 5caC was found enriched at low motif CTCF sites in primary T cells. CTCF and 5caC profiling in primary T cells and biochemical analysis support the notion that 5caC reinforces CTCF binding in suboptimal contexts. Together, these results provide a rationale for a perplexing aspect of CTCF biology: CTCF physically interacts with DNA methyltransferases that establish 5mC in genomic DNA and the TET proteins that site-specifically oxidize 5mC in a pathway to demethylation (Dubois-Chevalier et al., 2014; Guastafierro et al., 2008; Zampieri et al., 2012). Our results raise the intriguing possibility that CTCF association with these factors acts to reinforce its own binding, and potentially create a platform for the dynamic regulation of CTCF binding as DNMT and TET levels vary during development.

## RESULTS

### *De novo* CTCF sites overlap with 5caC-rich regions in *Tdg*-/- cells

Given the multiple critical roles played by CTCF in the nucleus, a deeper understanding of the molecular determinants that drive CTCF binding is warranted. We recently identified 5caC as one such factor, wherein we observed that purified CTCF showed increased interaction with 5caC-containing as compared to unmethylated DNA in electrophoretic mobility shift assays (EMSA) (Marina et al., 2016). However, whether 5caC enhances CTCF binding within the complex environment of chromosomal DNA was unclear. To begin to address CTCF/5caC binding *in vivo*, we turned to knockout mouse embryonic stem cells (mESCs) lacking the base-excision factor Tdg (*Tdg-/-*) (Fig. 1A) (Cortazar et al., 2011). Others previously established that Tdg depletion can be leveraged to boost the otherwise low level of 5caC within genomic DNA, without compromising other aspects of base-excision repair (Cortazar et al., 2011; He et al., 2011). Loss of Tdg expression in knockout mESCs was confirmed by immunoblotting (Fig. 1B) and a reciprocal increase in 5caC in *Tdg-/-* genomic DNA was demonstrated through dot blot with 5caC-specific antibody (Fig. 1C).

**Figure 1.**
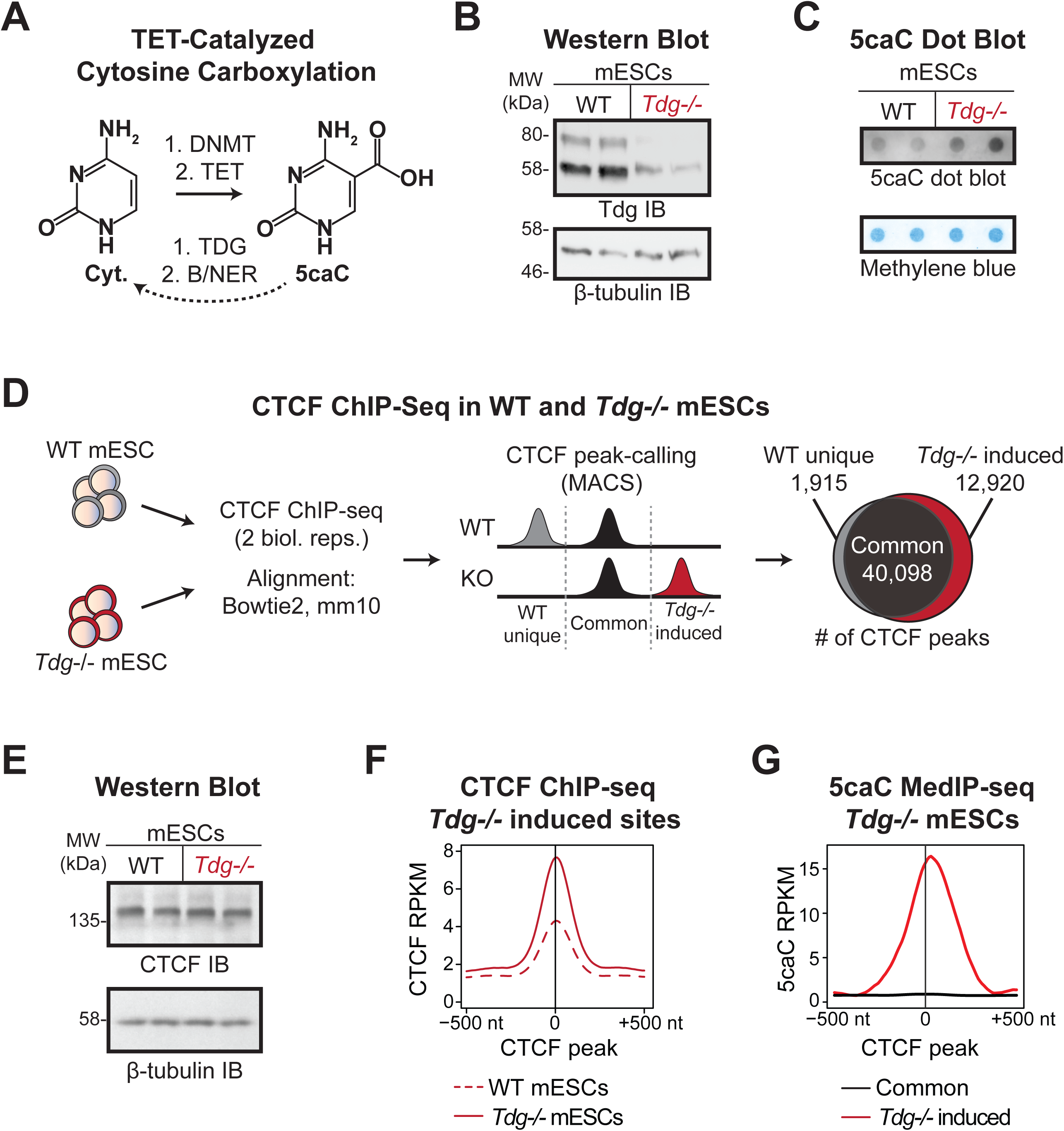
5caC enhances CTCF binding to genomic DNA. **(A)** Schematic of cytosine and 5-carboxylcytosine (5caC) molecular structures. **(B)** Tdg immunoblot in wildtype (WT) and *Tdg*-/- mESCs. Replicates are derived from independently maintained cultures. **(C)** 5caC dot blot in WT and *Tdg*-/- mESC genomic DNA. Replicates as in (B). **(D)** CTCF peaks determined through ChIP-seq in WT and *Tdg*-/- mESCs were parsed by occurrence in only WT or *Tdg-/-* cells (unique) or in both populations (common). The Venn diagram depicts the number of CTCF peaks per category. **(E)** CTCF immunoblot in WT and *Tdg*-/- mESCs. **(F)** CTCF ChIP-seq signal in WT and *Tdg*-/- mESCs centered on locations uniquely detected in *Tdg-/-* cells. Reduced CTCF signal in WT mESCs failed to reach the threshold for peak calling. **(G)** 5caC MedIP-seq in *Tdg*-/- mESCs centered on CTCF peaks unique to *Tdg-/-* cells or commonly detected in *Tdg-/-* and WT mESCs.

Having confirmed the elevated presence of 5caC in *Tdg-/-* cells, we next explored the consequence to CTCF binding genome-wide. CTCF chromatin immunoprecipitation and sequencing (ChIP-seq) was performed in duplicate in wildtype and *Tdg-/-* mESCs (Fig. 1D, Table S1). Consistent with previous reports, peak-calling indicated ∼40,000 CTCF peaks that were commonly detected in both mESC populations (Chen et al., 2012; The ENCODE Project Consortium, 2012; Wang et al., 2012). However, CTCF sites that were uniquely detected in one population or the other showed a dramatic imbalance: whereas ∼1,900 sites were present in wildtype but not *Tdg-/-* mESCs, nearly ∼13,000 sites were found in *Tdg-/-* but not wildtype mESCs (Fig. 1D, Table S2). In other words, Tdg loss was associated with a substantial increase in CTCF binding genome-wide. Importantly, this increase in CTCF binding was not related to a change in expression, as immunoblotting revealed comparable CTCF protein levels in wildtype and *Tdg-/-* mESCs (Fig. 1E). With this in mind, it is notable that comparison of CTCF ChIP-seq read density within the *Tdg-/-* induced sites showed that CTCF binding was not entirely absent in wildtype cells, but rather failed to reach the threshold for peak detection (Fig. 1F). These findings suggest that *de novo* CTCF binding in *Tdg-/-* cells results from a cellular change unrelated to CTCF abundance that reinforces binding at otherwise weak sites.

Based on our previous EMSA showing enhanced CTCF binding in the presence of 5caC, we specifically queried CTCF sites in *Tdg-/-* cells for emergent 5caC. To this end, we examined 5caC methylated DNA-immunoprecipitation sequencing (meDIP-seq) data from *Tdg*-depleted mESCs (Shen et al., 2013). Focusing within CTCF peaks, we observed a remarkable distinction in 5caC levels: while 5caC was absent from CTCF sites that were commonly detected in both wildtype and *Tdg-/-* mESCs, a robust 5caC signal was observed directly within *Tdg-/-* induced CTCF peaks (Fig. 1G). Together, these data raise the intriguing possibility that 5caC acts to reinforce CTCF binding in chromosomal DNA.

### *Tdg-/-* induced CTCF sites display unique molecular features *in vivo*

Having established the presence of 5caC-rich induced CTCF sites in *Tdg-/-* cells, we next explored their molecular basis and biological relevance. In particular, we examined for unique features in the induced subset as influenced by motif strength. Our rationale derived from two observations: 1) CTCF association with DNA does not adhere to a strict consensus sequence (Nakahashi et al., 2013; Rhee and Pugh, 2011) and 2) weak CTCF binding was observed in wildtype mESCs at locations that were robustly detected in *Tdg-/-* cells (Fig. 1F). These findings reflect a nuanced aspect of CTCF biology: the 11 zinc fingers associate with DNA to varying extents, resulting in degrees of binding strength and a relatively degenerate motif (Hashimoto et al., 2017; Nakahashi et al., 2013; Rhee and Pugh, 2011). We thus reasoned that overlapping 5caC at *Tdg-/-* induced CTCF sites may reflect a positive role for the cytidine modification in suboptimal contexts. Empirically determined CTCF binding sites in wildtype and *Tdg-/-* mESCs were accordingly assigned motif scores through the CTCFBSDB 2.0 database (Fig. 2A, schematic) (Ziebarth et al., 2013). Considering that 5caC was globally enriched in *Tdg-/-* induced CTCF sites (Fig. 1G), it is notable that the *de novo* sites were generally characterized by reduced motif strength as compared to commonly detected peaks (Fig. 2B). To specifically interrogate the relationship between 5caC and motif strength, CTCF peaks were segregated into groups representing low, mid and high motif scores (bottom quartile, middle quartiles, and top quartile, respectively) (Fig. 2A). CTCF sites with effectively no matching motif were represented in the low motif group. Examination of 5caC content within CTCF sites confirmed a general enrichment for 5caC across the spectrum of *Tdg-/-* induced sites, but also revealed an inverse relationship to CTCF motif strength, wherein 5caC levels were highest in the low motif subset (Fig. 2C). As 5caC is most frequently observed within CpG dinucleotide contexts, we further examined CpG content within CTCF sites parsed by motif strength. In agreement with increased substrate density, low motif sites were characterized by higher overall CpG density as compared to mid or high motif locations, thus providing a rationale for the increased prevalence of 5caC (Fig. 2D). These findings support a role for 5caC in promoting new CTCF binding in *Tdg-/-* cells, particularly to suboptimal motifs.

**Figure 2.**
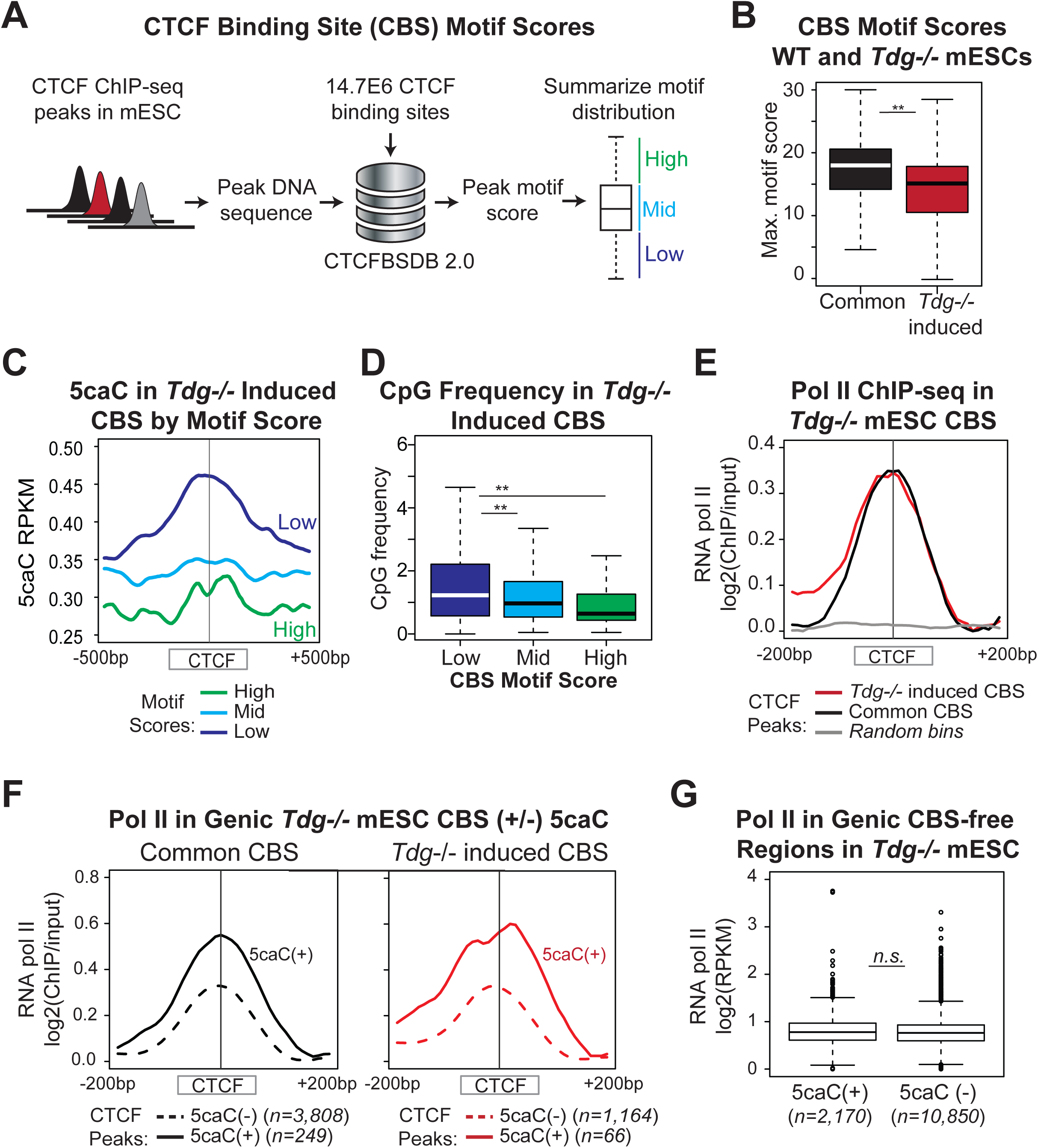
Low-motif CTCF peaks exhibit greatest gains in binding at 5caC-rich DNA. **(A)** CTCF ChIP-seq peaks in WT and *Tdg-/-* mESCs were assigned motif scores through the CTCFBSDB 2.0 algorithm and segregated into high, mid and low motif bins. **(B)** Box and whisker diagram of maximum motif score distribution in common vs. *Tdg-/-* induced CTCF peaks. **(C)** *Tdg-/-* mESC 5caC meDIP-seq within empirically determined CTCF peaks, segregated by motif score. **(D)** CpG dinucleotide frequency within *Tdg-/-* mESC CTCF peaks, segregated by motif score. **(E)** *Tdg-/-* mESC RNA pol II ChIP-seq density within common and *Tdg-/-* induced CTCF peaks relative to randomly permutated (shuffled) control peaks. **(F)** *Tdg-/-* mESC Pol II ChIP-seq in common and *Tdg-/-* induced CTCF sites as in (E), segregated on the basis of overlapping 5caC. **(G)** *Tdg-/-* mESC pol II ChIP-seq at 5caC-rich (+) and poor (-) regions not associated with proximal CTCF binding.

The above establish the presence of 5caC-rich CTCF sites in *Tdg-/-* cells, but to confirm functionality, we examined for co-occurrence of RNA polymerase II (pol II). We and others have shown that CTCF binding within genic DNA transiently obstructs pol II elongation, resulting in a local accumulation of pol II at CTCF sites (Lu and Tang, 2012; Paredes et al., 2013; Shukla et al., 2011). Consistent with the general distribution of CTCF binding, alignment to genomic features showed that ∼45% of the *Tdg-/-* induced sites occur within gene bodies (Fig. S2A, Table S3). To assess pol II levels within the induced sites, we performed pol II ChIP-seq in *Tdg-/-* cells. General examination of pol II read density within both common and induced genic CTCF sites revealed a clear accumulation that is consistent with *bona fide* CTCF binding, that was absent in randomized regions (Fig. 2E). To further explore the relationship to 5caC, pol II occurrence was examined within genic *Tdg-/-* induced CTCF sites segregated on the basis of overlapping 5caC (5caC-rich (+) or 5caC-poor (-)). While accumulating pol II was detected in both classes of CTCF sites, levels were markedly elevated for the 5caC(+) sites (Fig. 2F). Although somewhat unexpected, this finding is consistent with our previous demonstration of increased CTCF binding to a 5caC-containing probe in EMSA assay as compared to unmodified probe (Marina et al., 2016). Thus, the observed increase in pol II density at 5caC-rich CTCF sites in *Tdg-/-* cells raises the possibility that 5caC both promotes and strengthens CTCF binding *in vivo*. Importantly, examination of pol II in 5caC-rich versus poor genic regions that are not marked by proximal CTCF showed no distinction, demonstrating that accumulating pol II is not a general feature of 5caC-rich DNA (Fig. 2G; Fig. S2B). Together, these results establish both the presence and functionality of 5caC-associated CTCF sites in *Tdg-/-* cells.

Of note, mESCs are qualitatively distinct from differentiated tissues on several accounts. Most relevant to the current analysis, ES cells are uniquely characterized by non-CpG methylation and overall higher levels of 5mC-oxidized derivatives (Guo et al., 2014; Huang et al., 2014; Ito et al., 2010; Ito et al., 2011; Koh et al., 2011; Lister et al., 2009; Ramsahoye et al., 2000). Thus, to examine the generality of our findings outside of mESCs, we turned to primary human lymphocytes. We previously co-detected 5caC and CTCF at pre-mRNA splicing associated regions in naïve CD4+ T lymphocytes (Marina et al., 2016). TET1 and TET2 expression was high in naïve CD4+ T cells, whereas levels decreased upon activation (Marina et al., 2016). Accordingly, we isolated naïve CD4+ T cells from peripheral blood for genome-wide analysis of CTCF and 5caC through ChIP-seq and MedIP-seq, respectively (Fig. 3A). As in the mESC analysis, experimentally determined CTCF sites were parsed based on motif strength into low, mid and high scoring groups (Fig. 3B). Concordant with the mESC results, low motif CTCF sites in naïve CD4+ T cells were marked by increased CpG density and elevated overlapping 5caC as compared to the mid and high scoring groups (Fig. 3C, 3D). Given that it is commonly accepted 5caC levels within genomic DNA are too low for basal detection, 5caC occurrence within CTCF sites in an unperturbed cellular setting is noteworthy and attests to the relevance of association. Indeed, analysis of pol II ChIP-seq performed in naïve CD4+ T cells revealed a clear enrichment for pol II within low motif CTCF sites, which also showed the highest levels of overlapping 5caC (Fig. 3E).

**Figure 3.**
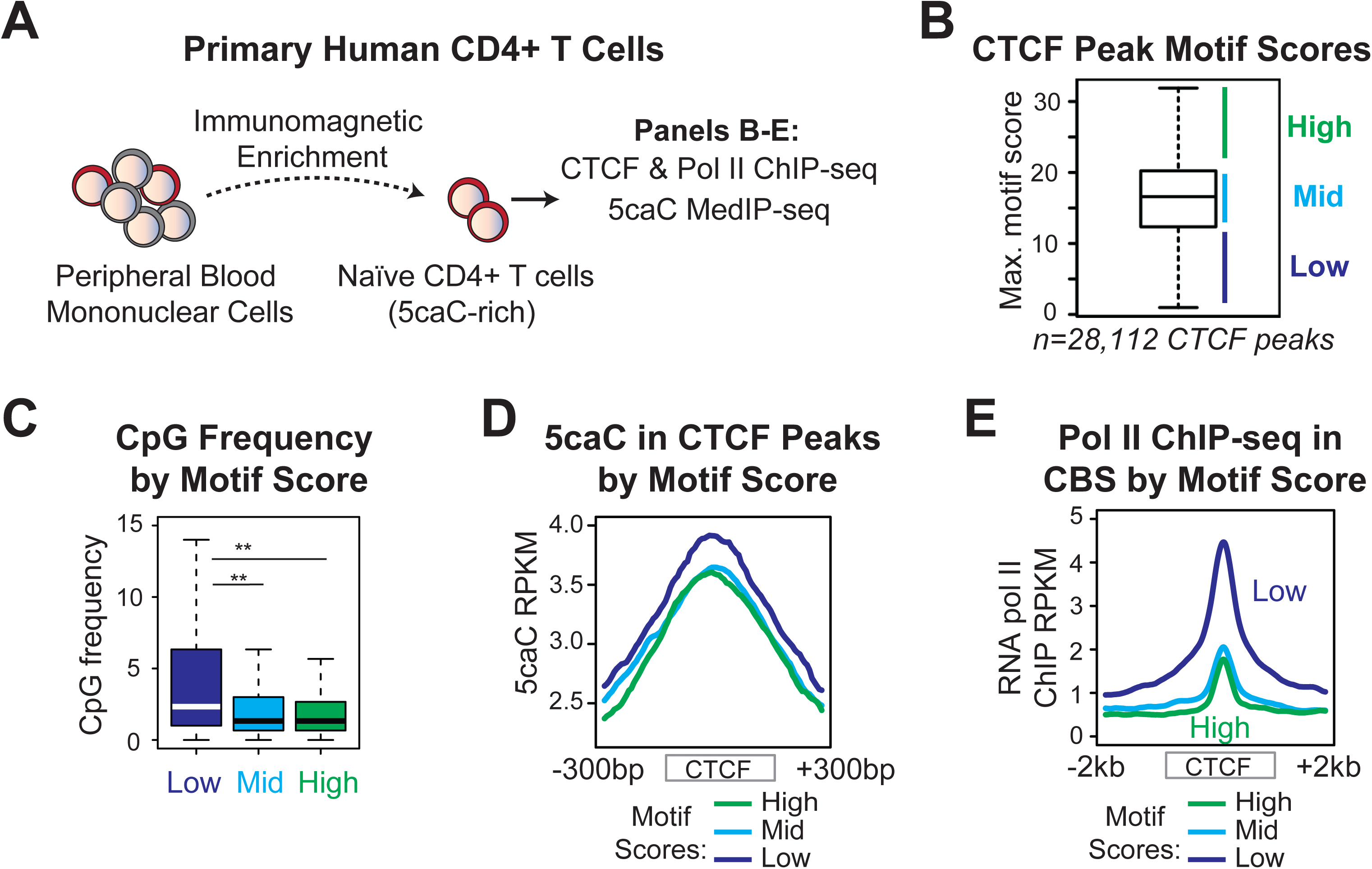
5caC associates with low motif CTCF binding sites in primary human CD4+ T cells. **(A)** Naïve CD4+ T cells were isolated from human peripheral blood through immunomagnetic enrichment and subjected to genome-wide analyses. **(B)** Distribution of maximum motif scores at CTCF peaks detected in naïve T cells. Peaks were segregated into high, mid, and low bins corresponding to scores within the top quartile, middle quartiles, and bottom quartile, respectively. **(C)** pG dinucleotide frequency within CD4+ T CTCF peaks, segregated by motif score. **(D-E)** 5caC meDIP-seq (D) and RNA pol II ChIP-seq (E) within determined CD4+ T CTCF peaks, segregated by motif score.

### CTCF binds motif-free DNA in the presence of 5caC *in vitro*

Our genome-wide data strongly support a role for 5caC in driving CTCF binding in a chromosomal setting. However, to move beyond correlations, we turned to *in vitro* systems for biochemical characterization of CTCF association with 5caC-containing DNA. In particular, based on the observed association of CTCF with 5caC-rich DNA at low motif sites, we examined whether 5caC promotes CTCF binding in suboptimal contexts through EMSA assays. Although EMSA varies from chromosomal DNA in the use of linear DNA probes, concerns related to artificiality are mitigated by the fact that CTCF binds to nucleosome-free DNA *in vivo* (Chen et al., 2012; Fu et al., 2008; Magbanua et al., 2015; Teif et al., 2014). Importantly, we previously revealed a surprising preference for 5caC-containing as compared to unmethylated DNA in CTCF EMSA: overall complex formation was increased in the presence of overlapping 5caC, and unlabeled 5caC was a more effective competitor of both complexes as compared to unmethylated competitor (Marina et al., 2016). These results were observed with multiple probes and distinct CTCF protein sources including recombinant CTCF and FLAG-tagged CTCF purified from cell culture (Marina et al., 2016). These findings are consistent with the genome-wide analysis of CTCF binding in *Tdg-/-* cells and highlight EMSA as an appropriate method for examining CTCF interaction with carboxylated DNA *in vitro*.

To validate the relationship between 5caC and CTCF motif strength uncovered in the genome-wide analysis, we generated EMSA probes embodying distinct CTCF binding modalities in the presence and absence of 5caC. To represent “strong” and “weak” motifs, respectively, we utilized a CTCF binding site located within exon 5 of the *CD45* gene in either a wildtype or mutated context. We previously performed extensive characterization of the *CD45* probes and showed that the incorporation of point mutations within the CTCF core to generate the weak probe abolished CTCF binding in EMSA with unmodified DNA (Marina et al., 2016; Shukla et al., 2011). Importantly, the introduced mutations do not alter CpG or general cytosine density in double-stranded DNA. For reference, the CTCFBSDB 2.0 algorithm revealed a score of 18.16 associated with the strong CTCF binding site, whereas the weak probe produced a score of 7.26 (Fig. 4A). EMSA was thus performed with FLAG-tagged CTCF purified from HEK293T lysates and radiolabeled *CD45* probes. 5caC incorporation was accomplished through: 1) PCR amplification in the presence of 2’-deoxy-5-carboxycytidine 5’-triphosphatase (5-carboxy-dCTP) or dCTP and 2) commercial synthesis. The PCR approach results in a 72 base pair probe centered on the CTCF binding site with uniformly modified cytosines, whereas commercial synthesis yields a 41mer with a total of 3 modified cytosines per DNA stand, occurring exclusively in a CpG context (Fig. 4A). Although CTCF binding is generally compromised on shorter probes such as the 41mer (Marina et al., 2016) both approaches yielded consistent and clear results. In the presence of unmethylated DNA, CTCF bound to the strong, but not the weak site, as evidenced through complex formation in phosphorimager analysis (Fig. 4B). Percent shift was calculated from phosphorimager analysis as the amount of label in complex with CTCF relative to free label (Fig. 4B). In contrast, CTCF formed a robust and specific complex with both the strong and weak probes in the presence of overlapping 5caC, as demonstrated through cold competition and supershift with anti-CTCF antibody (Fig. 4B).

**Figure 4.**
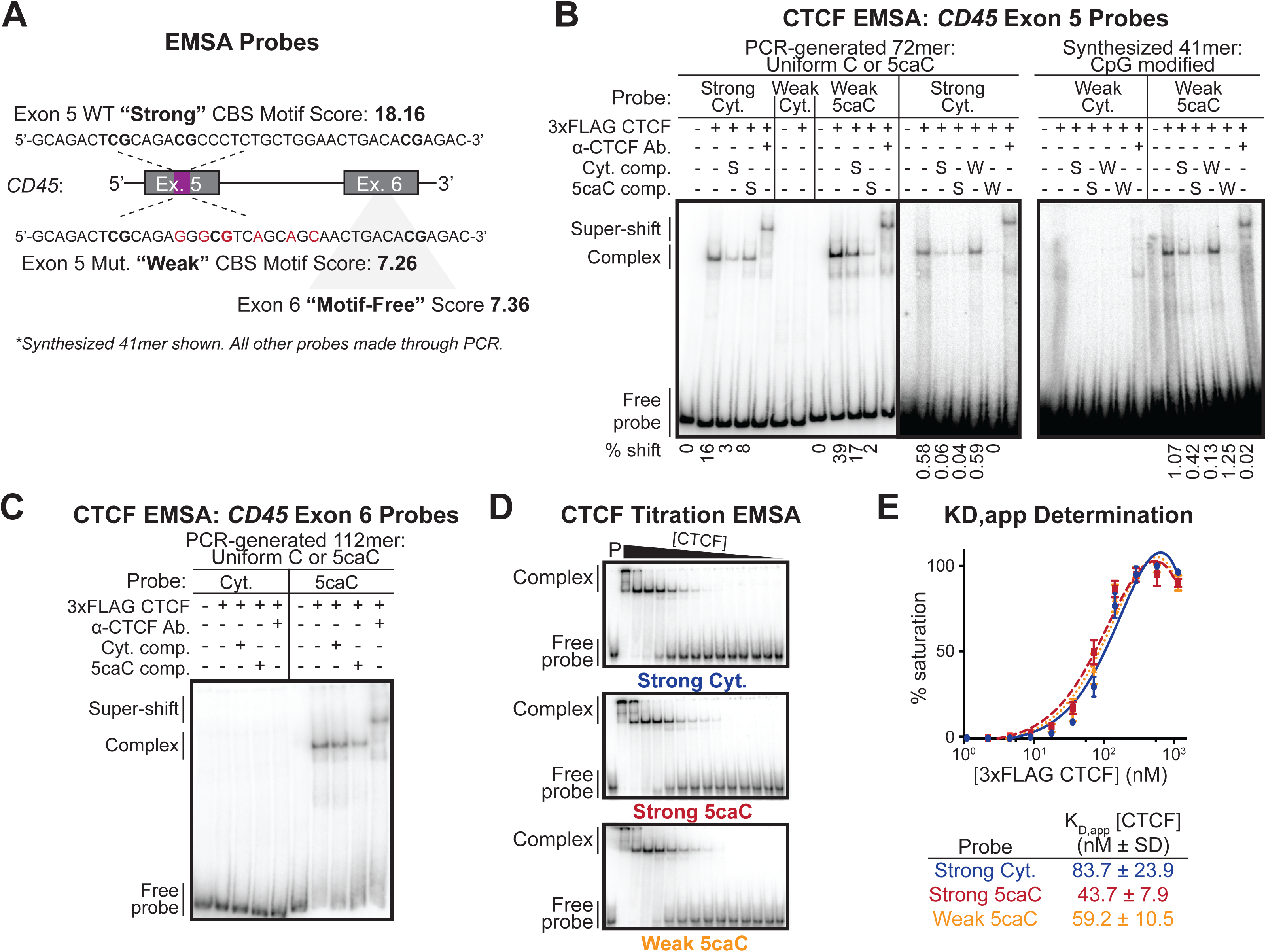
5caC enhances CTCF association with weak binding sites *in vitro*. **(A)** Schematic of EMSA probes derived from the human *CD45* gene. Point mutations were incorporated into a “strong” CBS within exon 5 to generate an analogous “weak” CBS probe. A region devoid of CTCF binding *in vivo* within exon 6 was selected as a “motif-free” control. CTCFBSDB 2.0 motif scores are shown. Bold sequences within the exon 5 probes show locations of CpGs and red text indicates mutated residues. **(B)** EMSA with affinity-purified CTCF and radiolabeled PCR-generated 72mer or commercially-synthesized 41mer DNA probes. Unlabeled strong (S) or weak (W) motif cold-competitor DNA added to indicated lanes; 10x and 100x molar excess used for 72mer and 41mer, respectively. Super-shift performed with α-CTCF antibody. Quantification of bound probe indicated as % shift below figure. **(C)** EMSA using affinity-purified CTCF and motif-free PCR-generated *CD45* exon 6 probes of varying carboxylation status; 10x molar excess of unlabeled CD45 exon 5 72mer used as cold-competitor where indicated. **(D)** Titration EMSA performed using 41mer *CD45* exon 5 DNA probes (3.76 nM); P indicates lane containing probe alone, CTCF concentration: 342.3-0.0418 nM. E) Saturation binding curves, derived from EMSAs in (D), and summary chart of apparent KD values for each 41mer probe; markers on curves indicate mean of two (Strong Cyt.) or three (Strong 5caC, Weak 5caC) independent replicates and error bars indicate standard deviation.

Of note, the genome-wide analysis of *Tdg-/-* induced CTCF sites revealed sequences for which the likelihood of CTCF binding was less than expected in random sequence (negative motif scores). Likewise, an unbiased mass spectrometry study that identified CTCF as a 5caC-specific reader utilized a probe entirely lacking elements consistent with known CTCF-interacting sequences (−8.04 motif score) (Spruijt et al., 2013; Ziebarth et al., 2013). To examine whether we could recapitulate CTCF-binding to seemingly motif-free DNA in the presence of 5caC, we performed additional EMSA with a sequence lacking any characteristics of CTCF binding. Specifically, *CD45* exon 6 does not contain any computationally predicted CTCF binding sites (7.36 motif score, Fig. 4A) and shows no evidence of CTCF binding in ChIP-qPCR and ChIP-seq (Marina et al., 2016; Shukla et al., 2011; The ENCODE Project Consortium, 2012). Exon 6 EMSA probes were generated through PCR in the presence of dCTP or 5-carboxy-dCTP. As predicted, purified CTCF did not interact with the unmethylated exon 6 probe under established binding conditions. In contrast, the 5caC-containing exon 6 probe formed a robust and specific complex with CTCF (Fig. 4C). This unexpected finding clearly demonstrates the positive impact of 5caC on CTCF binding *in vitro.*

The above EMSA establish that CTCF binding is qualitatively enhanced in the presence of 5caC. To further quantify the strength of association, we pursued relative binding affinity determination (K_D,apparent(app)_) through saturation binding experiments involving purified CTCF and radiolabeled *CD45* exon 5 41mers. EMSA was performed with a fixed amount of *CD45* probe representing strong (+/- CpG 5caC) and weak (+ CpG 5caC) CTCF sites and decreasing levels of purified CTCF (Fig. 4D) (Heffler et al., 2012). The unmodified weak probe was excluded from this analysis as CTCF binding was not detected in standard EMSA. Saturation binding curves were generated through percent shift to determine relative CTCF binding affinity as it relates to motif strength and 5caC (Heffler et al., 2012). Consistent with the enhanced binding visualized in standard EMSA, incorporation of 5caC into the three CpGs in the strong probe strengthened CTCF binding and resulted in a near 2-fold increase in affinity as compared to the unmodified strong binding site probe. Remarkably, the presence of 5caC within the weak motif probe yielded an intermediate K_D,app_ that was strengthened as compared to the unmodified strong motif probe but reduced relative to the 5caC-containing strong motif probe (Fig. 4E). These K_D,app_ values are within the established range for CTCF association with DNA (Hashimoto et al., 2017; Li et al., 2017; Martinez et al., 2014; Plasschaert et al., 2014). Taken together, these data corroborate the *in vivo* observations that overlapping 5caC, even in a minimal CpG setting, is sufficient to promote CTCF binding to weak consensus motifs.

An inherent limitation of EMSA relates to the fact that protein:DNA interactions are analyzed in the absence of additional variables at the cellular level. For example, while CTCF may interact with weak binding sites in the presence of 5caC in isolation, other factors may occupy such sites *in vivo*. To address this possibility, the strong and weak exon 5 probes were used as bait in DNA affinity purification assay (DAPA) (Fig. 5A). PCR-generated 72mer probes were biotinylated and immobilized on streptavidin-coated magnetic beads. Of note, capture of the 5caC-containing probes was slightly reduced relative to the unmodified probes, as assessed through SYBR Gold staining of the unbound portion (Fig. 5B). Nevertheless, recovery of weak versus strong probes was comparable per modification state, allowing for direct comparisons as related to motif strength (Fig. 5B). Incubation with nuclear extracts from HEK293T cells expressing FLAG-CTCF allowed for the capture and subsequent elution of associated proteins. Immunoblotting of DAPA eluates from the unmodified DNA probes demonstrated a robust interaction between CTCF and the probe containing a strong binding motif, whereas binding was not observed for the weak motif (Fig. 5C). In contrast, CTCF was recovered through incubation with both the strong and weak 5caC-containing probes. Consistent with the determined K_D,app_ CTCF association was reduced for the weak 5caC-containing probe, but was clearly visible (Fig. 5C). Relatedly, while the uneven streptavidin immobilization precludes direct comparison between unmodified and 5caC-containing probes, the overall reduction in CTCF recovery through the 5caC-containing probes may reflect competition for binding with other factors that shape the binding landscape *in vivo*. Overall, these DAPA results confirm that CTCF interacts with suboptimal DNA motifs in the presence of 5caC within the complex cellular milieu.

**Figure 5.**
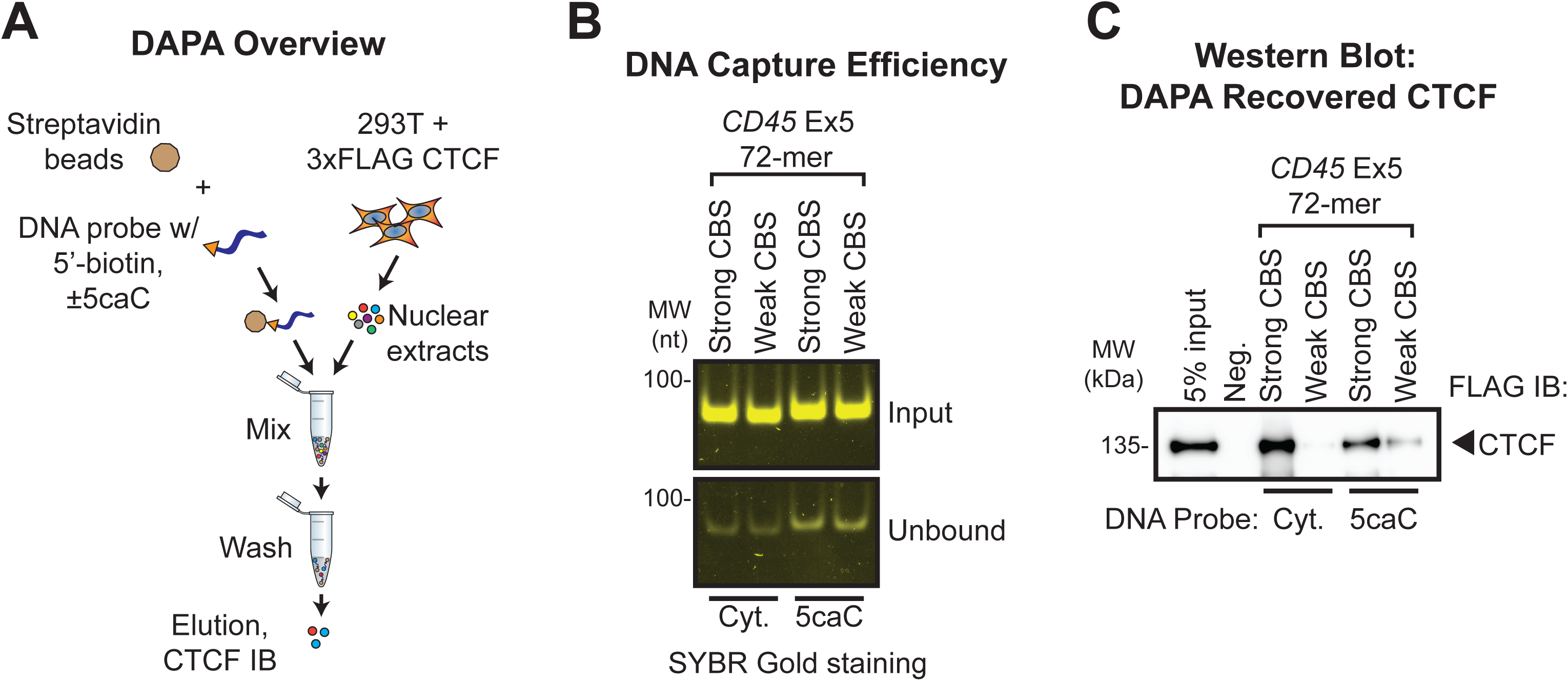
Affinity purification of CTCF from nuclear lysates with 5caC-containing DNA. A) Overview of CTCF DNA affinity precipitation assay (DAPA). Bead-bound DNA probes generated in the presence or absence of 5caC were used to enrich associated proteins from nuclear lysates. Probes representing strong and weak CTCF binding sites were used. SYBR Gold-staining to assess the capture efficiency of each DNA probe on streptavidin magnetic beads. C) Immunoblotting of DAPA-recovered CTCF through the various probes.

### CTCF generates a strong footprint at weak binding motifs in the presence of 5caC

The sum of the genome-wide analysis of *Tdg-/-* cells and general EMSA results clearly demonstrate that 5caC enhances CTCF binding in suboptimal contexts. However, to establish whether CTCF occupies a similar or unique expanse of DNA in the presence of 5caC, we pursued *in vitro* DNase I footprinting analysis. DNase I footprinting relies on time and concentration dependent DNase I cleavage of DNA according to availability (Brenowitz et al., 2001; Carey et al., 2013; Hampshire et al., 2007; Leblanc and Moss, 2015). Incubation of end-labeled probe with a protein of interest in the presence of DNase I can thus inform on the protein binding site through the region that is protected from cleavage (Brenowitz et al., 2001; Carey et al., 2013; Hampshire et al., 2007; Leblanc and Moss, 2015) (Fig. 6A). It is well-established that CTCF binds to nucleosome-free DNA and previous studies have demonstrated CTCF footprints of >20 bp (Chen et al., 2012; Filippova et al., 2001; Fu et al., 2008; Magbanua et al., 2015; Teif et al., 2014). We thus pursued DNase I footprinting with the synthesized *CD45* exon 5 41mer probe in which 5caC is restricted to 3 CpGs located within the CTCF binding core. Gel analysis of DNase I footprinting utilizing the unmodified strong motif probe in the presence of CTCF protein revealed a protected region encompassing the known location of CTCF binding (Figure 6B). In contrast, the unmodified weak CTCF motif probe failed to generate a clear protein-associated DNase I footprint (Fig. 6B). Although the weak motif probe displayed some protection from cleavage with increasing protein concentration, overall digestion patterns were virtually indistinguishable in lane histogram densitometry analysis in the presence or absence of CTCF (Fig. 6B). With this in mind, it is remarkable that the 5caC-containing weak motif generated a CTCF footprint that effectively mirrored the strong unmodified probe (Fig. 6B, 6C). These results solidify the role of 5caC in strengthening CTCF binding at weak binding sites. Curiously, however, the 5caC-containing strong motif probe did not generate a clear DNase I footprint (Fig. 6B). While the significance of this result is unclear, discrete regions of protection were observed in the lane histogram analysis (Fig. 6C). Taken into consideration with the K_D,app_ determination utilizing the same 41mer probes, overall attenuated digestion of the 5caC-containing strong motif probe may reflect a reduction in DNase I accessibility in response to particularly strong CTCF binding on the relatively short probe. Altogether, these DNase I footprinting data complement the genome-wide demonstration that 5caC enhances CTCF binding, particularly at suboptimal binding contexts.

**Figure 6.**
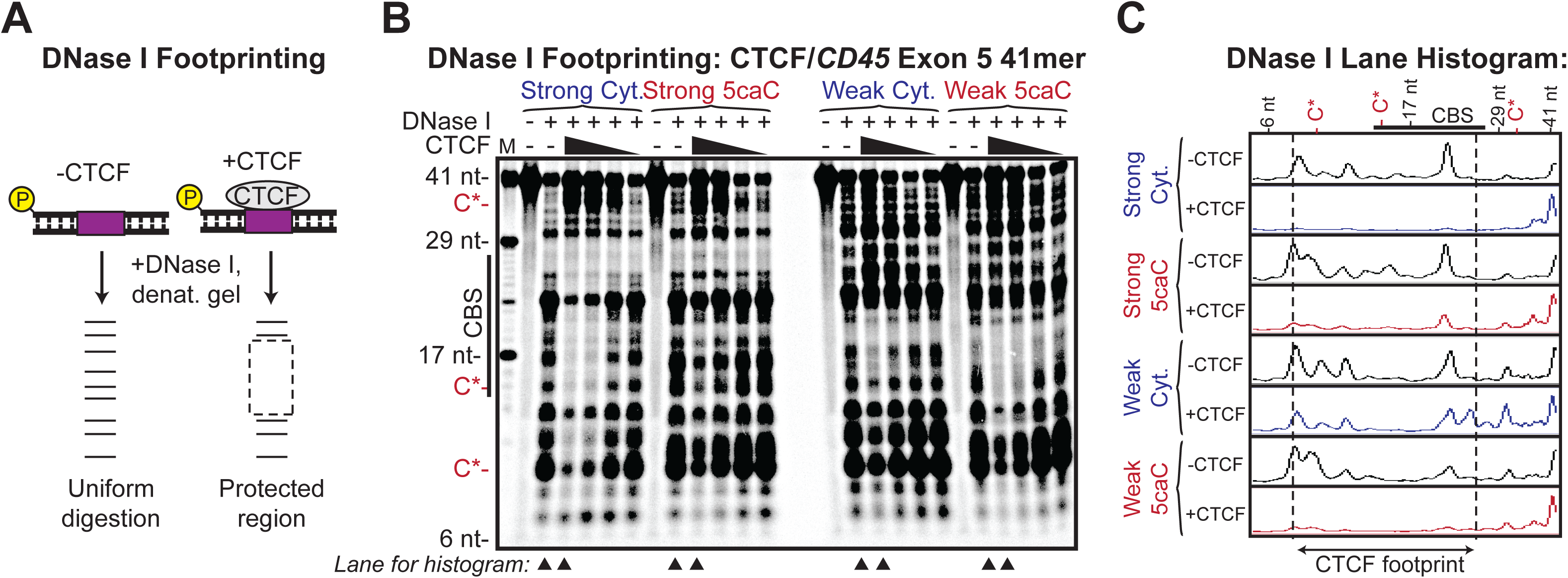
DNase I footprinting of CTCF associated with 5caC-containing DNA. **(A)** Overview of CTCF DNase I footprinting with variably carboxylated radiolabeled DNA probes representing strong and weak CTCF motifs. **(B)** Gel analysis of CTCF DNase I footprinting performed with the *CD45* exon 5 probe, ±CpG 5caC. The DNase I reaction contained 342.3-42.8 nM CTCF and 7.52 nM DNA probe. The location of carboxylated cytosine residues are indicated by C*. M signifies oligonucleotide marker and arrowheads indicate lanes used to generate histograms. **(C)** Lane histogram densitometry analysis of the indicated lanes from (B). The region protected by CTCF binding is shown.

## DISCUSSION

Once considered a stable hereditary mark, it is now appreciated that DNA methylation is dynamically regulated to shape and define gene expression in a cell-specific manner (reviewed in Luo *et al.*, 2018) (Luo et al., 2018). Tissue-specific changes in methylation often occur within gene bodies (Deaton et al., 2011; Maunakea et al., 2010), wherein we previously determined a role in pre-mRNA splicing that is achieved through modulation of CTCF binding (Marina et al., 2016; Shukla et al., 2011). We showed that genic CTCF promotes inclusion of weak exons in spliced mRNA through local RNA polymerase II pausing, whereas DNA methylation has the opposite effect (Shukla et al., 2011). Our efforts to understand how dynamic methylation at CTCF sites is achieved further uncovered a role for TET-catalyzed oxidized 5mC derivatives (5oxiC) (Marina et al., 2016). In particular, we asked how methylation is modulated at specific CTCF sites while leaving others unaffected. Given that CTCF is critical to numerous cellular processes, including general nuclear architecture, precise control of variable binding would be of tantamount relevance (Dixon et al., 2012; Handoko et al., 2011; Rao et al., 2014; Vietri Rudan et al., 2015; Zuin et al., 2014). We previously determined that splicing-associated ‘dynamic’ CTCF sites are marked by overlapping oxidized derivatives, whereas static sites are unmethylated (Marina et al., 2016). Considering that CTCF has been biochemically associated with both the DNMT enzymes (that would evict CTCF) (Guastafierro et al., 2008; Zampieri et al., 2012) and the TET enzymes (that would facilitate CTCF binding) (Dubois-Chevalier et al., 2014) these results raised the possibility that 5oxiC facilitates CTCF binding and bookmarks locations of future CTCF eviction as TET activity declines. In other words, CTCF association with DNMT1 would ensure methylation of proximal CpGs post-replication, whereas the TET enzymes would subsequently oxidize 5mC and enable CTCF binding. However, a problematic aspect of this model related to the fact that CTCF is incapable of binding the abundant 5mC oxidized derivative 5hmC (Marina et al., 2016), and the downstream derivatives 5fC and 5caC are lowly detected in genomic DNA (Ito et al., 2011; Lu et al., 2015).

Our demonstrations associating CTCF and 5caC reconcile these observations: CTCF robustly interacts with 5caC, *in vitro* and *in vivo*, and 5caC is readily detected at CTCF sites. This latter point further suggests that CTCF may protect 5caC from removal through the base excision repair enzymes (He et al., 2011). However, the truly unexpected aspect of this work relates to the observation that CTCF binding is seemingly enhanced through overlapping 5caC, suggesting a novel mode of binding. These results are consistent with a previous unbiased mass spectrometry study employing a short CpG-carboxylated DNA probe, wherein CTCF was identified as a 5caC specific reader (Spruijt et al., 2013). As in our EMSA experiments, this DNA fragment lacked a computationally-identifiable CTCF binding site. Combined with the genome-wide observation that CTCF binding sites with overlapping 5caC are generally characterized by lower motif scores, these findings suggest a unique mode of binding. While the precise biophysical determinants guiding CTCF binding at unmethylated versus 5caC rich DNA are unclear, our DNase I footprinting data comparing an unmodified strong versus carboxylated weak CTCF motif suggest that the binding modes are conformationally similar. The apparent increase in affinity in the presence of 5caC may thus relate to charge-based stabilization of CTCF zinc-finger contacts. Alternatively, the presence of 5caC within genomic DNA may facilitate a double-helical structure that enables CTCF binding. Detailed biophysical studies involving CTCF and 5caC-containing DNA will be required to resolve these points.

Notably, a recent GRO-seq study described a role for 5caC in reduced RNA polymerase II elongation in *Tdg-/-* cells (Wang et al., 2015). Our description of ∼13,000 novel CTCF sites upon *Tdg* deletion raises the possibility that emergent CTCF contributed to the overall reduction in pol II processivity in *Tdg-/-* cells. Indeed, in our hands, examination of pol II occupancy at 5caC rich regions showed no elevation as compared to 5caC poor regions. Further analyses will be required to conclusively determine the source of altered elongation in *Tdg-/-* cells.

In sum, we describe, for the first time, a role for 5caC in modulating CTCF binding in cells. Given that 5caC levels vary during development (Wheldon et al., 2014), these results have significant implications to dynamic CTCF binding during tissue differentiation. A detailed analysis of CTCF and 5caC co-occurrence during organismal development will inform on the extent to which 5caC shapes CTCF tissue specificity. Importantly, our findings reported herein raise the possibility that CTCF sites may be engineered in the genome through targeted 5caC. As CRISPR/Cas9 technology continues to advance, one can envision applications both in studying specific CTCF sites and the overall impact of induced CTCF on nuclear architecture.

## Supporting information

## Acknowledgements

We thank the members of the Center for Cancer Research Sequencing Facility (CCR-SF) at the National Cancer Institute (Frederick, MD) for providing Illumina sequencing services. We thank Dr. Primo Schär of the University of Basel for providing wildtype and *Tdg-/-* mESCs. This study utilized the Biowulf Linux cluster at the National Institutes of Health, Bethesda, MD (http://biowulf.nih.gov). This work is supported by the Intramural Research Program of NIH, the National Cancer Institute, The Center for Cancer Research.

## Author Contributions

Conceptualization, S.O.; Methodology, K.K.N., M. T. and S.O.; Data analysis and curation, D.M.S. and A.A.D.; Investigation and validation, K.K.N., M.F.P., A.A.D and M.D.M.; Writing, K.K.N, D.M.S. and S.O.; Supervision and funding acquisition, S.O.

## Supplementary Figure Titles and Legends

**Figure S2.**
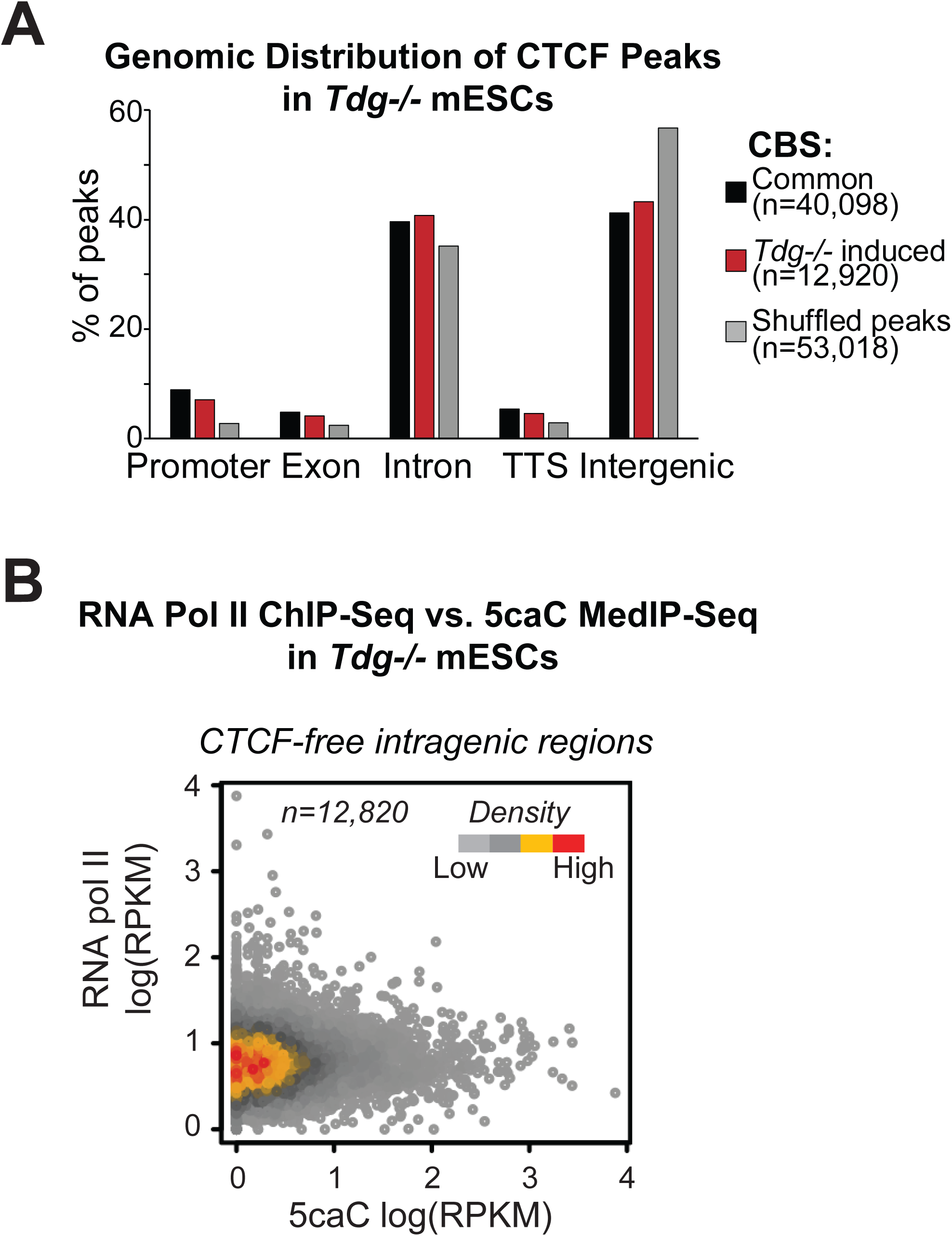
related to Figure 2. Genome-wide analysis of CTCF, 5caC and RNA pol II in *Tdg-/-* mESCs. A) Genomic distribution of common and *Tdg-/-* induced CTCF ChIP-seq peaks throughout the murine genome, compared to randomly permutated (shuffled) peaks. B) Scatter plot comparison of RNA pol II ChIP-seq and 5caC MedIP-seq densities at CTCF-free intragenic regions in *Tdg-/-* mESC. Increasing 5caC levels do not correlate with increased pol II.

## METHODS

### CONTACT FOR REAGENT AND RESOURCE SHARING

Further information and requests for resources and reagents should be directed to and will be fulfilled by the Lead Contact, Shalini Oberdoerffer. There are no restrictions on any data or materials presented in this paper.

## EXPERIMENTAL MODEL AND SUBJECT DETAILS

### Cell culture and transfection

293T acquired from the ATCC were cultured in DMEM (Gibco) supplemented with 2 mM L-glutamine (Gibco) and 10% bovine calf serum (HyClone, #SH30073.03). *Tdg*-/- mouse embryonic stem cells (mESCs) and wildtype control mESCs were a gift from Dr. Primo Schär. Complete mESC media was composed of KnockOut DMEM (Gibco), supplemented with 2 mM GlutaMAX (Gibco), non-essential amino acids (Gibco), 0.1 mM 2-mercaptoethanol (SIGMA), 1,000 U/ml LIF (Millipore, #ESG1106), and 15% ES-tested FBS (Gibco, #10439-024). mESC were co-cultured with mitotically-arrested mouse embryonic fibroblast (MEF; produced in-house or purchased from Millipore, #PMEF-CF) on gelatin-coated (Millipore, #ES-006-B) dishes. MEF depletion was achieved by three 10-minute rounds of serial-plating onto uncoated tissue culture dishes. Human CD4+ human T lymphocytes were acquired and cultured as described in Marina *et al.*, 2016. Plasmids were transfected into 293T using Lipofectamine 2000 (Invitrogen, #11668019) according to the manufacturer’s instructions.

## METHOD DETAILS

### Antibodies

The following antibodies were used for immunoblotting: anti-β-tubulin (Cell Signaling Technology; CST, #2146), anti-CTCF (CST, #3418), anti-FLAG (Sigma, #F3165), anti-TDG (abcam, #ab154192), HRP-linked anti-mouse IgG (Jackson ImmunoResearch, #115-035-003), HRP-linked anti-rabbit IgG (CST, #7074). The following antibodies were used for ChIP: anti-CTCF (CST, #3418), anti-RNA pol II (Millipore, #05-623), monoclonal rabbit isotype control IgG (DA1E) (CST, #3900), normal rabbit polyclonal control IgG (CST, #2729). Anti-5caC (abcam, #ab185492) was used for DNA dot blot.

### 5caC dot blot

Genomic DNA was purified from MEF-depleted *Tdg*-/- and control mESC using PureLink Genomic DNA Mini Kit (Invitrogen, #K182001) according to the manufacturer’s instructions. Dot blot was performed as described by the Cell Signaling Technology protocol for product #36836. Briefly, genomic DNA was sheared between 200 and 500 bp by sonication (Bioruptor Twin, Diagenode). 10 µg fragmented DNA was heated at 95 °C for 10 minutes in 200 µl DNA denaturing buffer (100 mM NaOH, 10 mM EDTA), neutralized with 200 µl 20X SSC (3 M NaCl, 300 mM sodium citrate, pH 7.0), and chilled on ice for 5 minutes. DNA was applied to nylon membrane (GE Healthcare, #RPN303B) using a Bio-Dot apparatus (Bio-Rad), air-dried then fix to membrane by UV crosslinking (Stratalinker 2400, Stratagene). Membrane was blocked at room temperature for 1 hour in PBS containing 0.05% Tween-20 (PBS-T), 2.5% BSA, and 2.5% non-fat dry milk and probed overnight at 4 °C with 5caC antibody (abcam, #ab185492) diluted 1:1000 in blocking buffer. After washing and incubation with HRP-linked anti-rabbit 2° antibody diluted 1:1000 in blocking buffer, signal detection was performed using SuperSignal West Femto ECL substrate (Thermo Scientific, #34095), imaged using ChemiDoc MP (Bio-Rad), and densitometric quantification performed using ImageLab software (Bio-Rad). Genomic DNA samples were applied to a separate nylon membrane as described above and methylene blue was used as a total DNA stain that served as loading control.

### Chromatin immunoprecipitation-sequencing (ChIP-seq)

Prior to chromatin immunoprecipitation, antibodies (5 µl polyclonal or 5 µg monoclonal) were pre-bound to 200 µl Protein G magnetic beads (Invitrogen, #10004D) by overnight incubation in PBS with 5% BSA; beads were washed and resuspended in PBS with 5% BSA. MEF-depleted mESCs were fixed with 1% formaldehyde (Sigma-Aldrich, # 252549) at room temperature for 5 min and quenched with 125 mM glycine (ICN Biomedical, #ICN808822). Cell membranes were lysed with cold NP-40 buffer (1% NP40, 150 mM NaCl, 50 mM Tris–HCl; pH 8.0) and nuclei collected by centrifugation at 12,000 × g for 1 min at 4 °C. Nuclear pellets were resuspended at a concentration of 2E8 cells/ml in ChIP sonication buffer (1% SDS, 10 mM EDTA, 50 mM Tris– HCl; pH 8.0) supplemented with protease (Thermo Scientific, #78430) and phosphatase (Calbiochem, #524627) inhibitors. Chromatin was sheared into 150-400 bp fragments by sonication (Bioruptor Twin, Diagenode). Debris was pelleted by centrifugation and cleared chromatin was diluted 10-fold in ChIP dilution buffer (1.1% Triton X-100, 0.01% SDS, 167 mM NaCl, 1.2 mM EDTA, 16.7 mM Tris–HCl; pH 8.1). The prepared antibody-bound beads were added to 1 ml diluted chromatin containing 2E7 million cell equivalents and incubated overnight with rotation at 4 °C. Immune complexes were washed 5 times with LiCl wash buffer (250 mM LiCl, 1% NP-40, 1% sodium deoxycholate, 100 mM Tris-HCl; pH 7.5) and once with TE (0.1 mM EDTA, 10 mM Tris-HCl; 7.5). Beads were resuspended in IP Elution Buffer (1% SDS, 0.1 M NaHCO_3_) and crosslinks reversed by overnight incubation at 65 °C. DNA was purified by column purification (QIAGEN, # 28106) and subjected to Illumina sequencing at the Advanced Technology Research Facility at NCI Frederick. Libraries were constructed with the Illumina TruSeq ChIP Sample Prep Kit (#IP-202-1012-1024) and sequenced on an Illumina NextSeq instrument for seventy-six cycles in single-end mode, using NextSeq High Output v.2 chemistry.

### Nuclear extracts

Nuclear extracts were prepared from 293T cells, either untransfected or transfected with plasmid encoding 3xFLAG-hCTCF as described in Jacobs *et al.*, 1993 (Jacobs et al., 1993). Hypotonic lysis buffer A (20 mM HEPES; pH 7.9, 10 mM KCl, 1 mM EDTA, 1 mM DTT) and nuclear extraction buffer B (20 mM HEPES; pH 7.9, 400 mM NaCl, 1 mM EDTA, 1mM DTT) were supplemented with 4 µM ZnCl_2_, protease inhibitor (Thermo Scientific, #78429) and Calbiochem phosphatase inhibitors (EMD-Millipore, #524627). Protein quantification was performed using Bradford assay (Bio-Rad #5000201).

### DNA affinity precipitation assay (DAPA)

Double-stranded *CD45* exon 5 DNA probes were generated by amplification of plasmid DNA template with Phusion polymerase (NEB, #M0530L) using 5’ biotinylated forward (5’-CCT CAC CTT CCC ACG CAC GCA GAC TC-3’) and unlabeled reverse primers (5’-GGA GCC GCT GAA TGT CTG CGT GTC AGT TC-3’) (Integrated DNA Technologies). PCR performed in the presence of dCTP and 5-carboxy-dCTP (Trilink, #N-2063) produced unmodified or uniformly carboxylated DNA probes, respectively. Biotinylated probes were immobilized on streptavidin magnetic beads (Invitrogen, #11205D) according to the manufacturer’s instructions; DNA capture efficiency was evaluated by monitoring unbound DNA present in the supernatant by SYBR Gold (Invitrogen, # S11494) staining after gel electrophoresis. DAPA was performed by combining 20 µg nuclear extract with 50 mg DNA-bound streptavidin beads in 100 µl DAPA buffer (10 mM Tris-HCl; pH 8.5, 50 mM KCl, 0.1 mM ZnCl2, 0.1% NP-40, 0.05 mg/ml BSA, 2 mM DTT, 1 ug/ml poly-dI:dC) supplemented with HALT protease inhibitor (Thermo Scientific, #78429) and Calbiochem phosphatase inhibitor (EMD-Millipore, #524627) cocktails. After overnight incubation with rotation at 4 °C, beads were washed 4 times with 100 µl DAPA buffer, heated at 95 °C for 10 minutes in Laemmli sample buffer, and subjected to immunoblotting using standard techniques.

### CTCF purification, EMSA, and relative KD determination

CTCF was affinity-purified from 293T cells transfected with a 3xFLAG-tagged CTCF expression construct using FLAG M2 agarose beads as described in Marina *et al.*, 2016 with the exception that the NaCl concentration in lysis and wash buffers was increased to 500 mM. Purified CTCF was quantified relative to Precision Plus Protein standards (Bio-Rad, #1610363) in a Coomassie-stained gel imaged on Bio-Rad ChemiDoc MP and analyzed with Bio-Rad ImageLab software. *CD45* exon 5 72-mer was produced using PCR amplification as indicated above; *CD45* exon 6 was produced by PCR amplification of template plasmid DNA using unlabeled forward (5’-AGC ACC TTT CCT ACA GAC CCA GTT-3’) and reverse (5’TGT TCG CTG TGA TGG TGG TGT T-3’) primers. DNA probes were labeled using γ-^32^P-ATP (Perkin Elmer #NEG502A250UC) and PNK (NEB #M0201L) and EMSA was performed as described in Marina *et al.*, 2016; cold competition assays utilized 10x and 100x molar excess of unlabeled 72-mer and 41-mer DNA probes, respectively. Relative KD was determined using EMSA as described in Heffler et al., 2012 (Heffler et al., 2012). Briefly, CTCF EMSA was first performed by incubating two-fold serial dilutions of purified 3xFLAG-CTCF with a fixed amount of radiolabeled DNA probe. The fraction of CTCF-bound DNA was determined using background subtracted densitometric signals as follows; fraction bound = complex/(complex + free probe). Saturation binding curves were generated by plotting fraction bound relative to CTCF concentration and non-linear regression was used to obtain relative KD values (Prism 7, GraphPad). Data are represented as mean ± SD for at least 2 replicates.

### DNase I footprinting

The sense strand of each commercially-synthesized 41-nt *CD45* exon 5 probe was individually radiolabeled using γ-32P-ATP and PNK as described above and annealed to equimolar quantities of its unlabeled complementary strand in annealing buffer (10 mM Tris, pH 8.0, 50 mM NaCl, 1 mM EDTA) by heating to 95 °C and allowing to cool slowly to room temperature. Free nucleotides were separated from the annealed probe using the QIAquick Nucleotide Removal Kit (QIAGEN, #28304). Probes were quantified using Qubit dsDNA HS Assay Kit (Invitrogen, #Q32851). DNase I footprint was performed by combining CTCF and radiolabeled DNA fragment in CTCF DNase I footprinting buffer (10 mM Tris-HCl; pH 8.5, 0.5 mM CaCl2, 2.5 MgCl2, 50 mM KCl, 0.1 mM ZnCl2, 0.1% NP-40, 0.05 mg/ml BSA, 2 mM DTT, 1 ug/ml poly-dI:dC) and incubating on ice for 30 minutes. DNase I (1.25E-3U/ul; NEB, #M0303L) was added and samples were incubated at room temperature for the indicated time intervals. DNase I digestion was stopped by addition of equal volume of 2X formamide loading buffer (95%formamide, 20 mM EDTA, 0.025% each of xylene cyanol, bromophenol blue, orange G) and heating at 95 °C for 3 minutes before loading on a pre-run 15% 7M urea gel. Gel was exposed to phosphor screen (Molecular Dynamics) for 24 hours; signals were detected using PhosphorImager (Storm 840, Molecular Dynamics).

## Bioinformatics

### Genomic alignment, peak-calling, annotation, and accession numbers

ChIP-seq reads from our experiments and published data were aligned to the relevant reference genome (mm10 or hg19) using Bowtie version 2.2.9 (Langmead and Salzberg, 2012) with default parameters; alignment statistics are presented in Table S1. Peak calling was performed with MACS (v 2.2.1) (Zhang et al., 2008), using sonicated chromatin (input) as the control. Peaks were called separately for each replicate. Peaks called on chromosome Y were removed. Merged peak calls from replicates constituted the reference set, from which a set of consensus peaks was defined based on presence in individual replicates; all subsequent analyses were performed using these peak coordinates (Table S2). Peak annotation was determined for the mm10 genome using HOMER version 4.10 with default settings (Table S3); genomic annotation of an equal number of shuffled peaks was used to derive observed/expected values (Heinz et al., 2010). Human CD4+ T cell CTCF ChIP-seq data were previously reported in Marina *et al.*, 2016 (GEO accession #GSE74850) (Marina et al., 2016). 5caC meDIP from control and *Tdg*-depleted mESC were reported in Shen et al., 2013 (GEO accession #GSE46111) (Shen et al., 2013). CD4+ T pol II ChIP-Seq data were obtained from Barski et al., 2007 (Barski et al., 2007). CTCF and RNA pol II ChIP-seq data from *Tdg-/-* and wildtype control mESCs are accessible via GSEXXXX).

To generate aggregate plots from sequencing data, deepTools was used to compute coverage summaries from bigwig files (Ramirez et al., 2014). Bedtools was used for coordinate intersections (Quinlan and Hall, 2010). When intersecting features with transcript models to define intergenic/intragenic, we used Ensembl rel. 83 for mm10. To define an expressed gene in mESC we aligned reads from poly(A) RNA-seq performed in wildtype and *Tdg-/-* mESCs (GSEXXXX) with Tophat v2.1.1 (Trapnell et al., 2009) and quantified gene expression with Cufflinks 2.2.1 (Trapnell et al., 2010). Genes with fragments per kilobase per million mapped reads (FPKM) greater than 1 were considered expressed genes. Genic definitions for T-cell data were taken from Marina *et al.*, 2016 (Marina et al., 2016).

### Evaluation of CTCF consensus binding sites

CTCF binding sites were detected and scored using the “Scan” feature in CTCFBSDB 2.0 (Table S2) from peak sequence (Ziebarth et al., 2013). This tool takes user-input sequence and queries a set of curated Position Weight Matrices (PWMs) to generate match scores. Raw database results were imported into R for downstream filtering and analysis. Unless otherwise indicated, only the maximum motif score for each input sequence is reported. To calculate CpG frequencies, in house perl scripts were used to count CG frequency relative to all other dinucleotide combinations as described in Marina *et al.*, 2016 (Marina et al., 2016).

## Supplemental Item Titles

Table S1. Summary of CTCF ChIP-seq alignment statistics in mESCs, related to Fig. 1.

Table S2. Locations and properties of wildtype and *Tdg -/-* mESC CTCF peaks, related to Figs. 1 and 2. xlsx

Table S3. Genomic annotation of *Tdg -/-* mESC CTCF peaks using HOMER related to Fig. 2 and S2.xlsx

